# Human induced pluripotent stem cell-derived neuroectodermal epithelial cells mistaken for blood-brain barrier-forming endothelial cells

**DOI:** 10.1101/699173

**Authors:** Tyler M. Lu, David Redmond, Tarig Magdeldin, Duc-Huy T. Nguyen, Amanda Snead, Andrew Sproul, Jenny Xiang, Koji Shido, Howard A. Fine, Zev Rosenwaks, Shahin Rafii, Dritan Agalliu, Raphaël Lis

## Abstract

Brain microvascular endothelial cells (BMECs) possess unique properties underlying the blood-brain-barrier (BBB), that are crucial for homeostatic brain functions and interactions with the immune system. Modulation of BBB function is essential for treatment of neurological diseases and effective tumor targeting. Studies to-date have been hampered by the lack of physiological models using cultivated human BMECs that sustain BBB properties. Recently, differentiation of induced pluripotent stem cells (iPSCs) into cells with BBB-like properties has been reported, providing a robust *in vitro* model for drug screening and mechanistic understanding of neurological diseases. However, the precise identity of these iBMECs remains unclear. Employing single-cell RNA sequencing, bioinformatic analysis and immunofluorescence for several pathways, transcription factors (TFs), and surface markers, we examined the molecular and functional properties of iBMECs differentiated either in the absence or presence of retinoic acid. We found that iBMECs lack both endothelial-lineage genes and ETS TFs that are essential for the establishment and maintenance of EC identity. Moreover, iBMECs fail to respond to angiogenic stimuli and form lumenized vessels *in vivo*. We demonstrate that human iBMECs are not barrier-forming ECs but rather EpCAM^+^ neuroectodermal epithelial cells (NE-EpiCs) that form tight junctions resembling those present in BBB-forming BMECs. Finally, overexpression of ETS TFs (ETV2, FLI1, and ERG) reprograms NE-EpiCs to become more like the BBB-forming ECs. Thus, although directed differentiation of human iBMECs primarily gives rise to epithelial cells, overexpression of several ETS TFs can divert them toward a vascular BBB *in vitro*.

## Introduction

Brain microvascular endothelial cells (BMECs) which line the vascular network of the central nervous system (CNS) form a specialized barrier termed the blood-brain barrier (BBB) that regulates the dynamic transfer of select molecules into and out of the CNS. The BBB also restricts the entry of immune cell and is essential for healthy brain activity. These properties are achieved through the presence of specialized tight junctions exhibiting a high transendothelial electrical resistance (TEER) *in vivo*, reduced caveolar-mediated transport and the presence of selective transporters^1–3^. Loss of BBB integrity is a hallmark of many neurological diseases such as ischemic stroke and multiple sclerosis and contributes to tumor metastasis and pathogenesis of glioblastoma multiforme (GBM)^1,4^. The presence of a neurovascular barrier, on the other hand, impedes successful delivery of therapeutics into the CNS in many neurological and psychiatric diseases. The development of *in vitro* models has accelerated mechanical studies on the BBB as well as large scale screening of drugs with potential to penetrate the brain, with some limitations^5–9^. Human primary BMECs are difficult to procure in sufficient numbers for experimental purposes, and their barrier properties cannot be maintained *in vitro*. Many studies have used immortalized human BMECs, however these cells lose their BBB attributes and produce a sub-physiologic TEER, rendering them ineffective for functional studies^8^.

Recently, the generation of a pure population of putative BMECs from pluripotent stem cell sources (iBMECs) has been described to meet the need for a reliable and reproducible *in vitro* human BBB model^10–22^. Human pluripotent stem cells, embryonic or induced, can differentiate into large quantities of specialized cells in order to study development and model disease^23–27^. iBMECs are generated primarily through differentiation of pluripotent stem cells into neural and endothelial progenitors, followed by selective purification (referred to as neuroendothelial differentiation)^10,28,29^. Adding retinoic acid or inhibiting GSK3 during the differentiation process reportedly enhances the yield and fidelity of these putative iBMECs^10,30^. These iBMECs display endothelial-like properties, such as tube formation and low-density lipoprotein uptake, high transendothelial electrical resistance (TEER) and barrier-like efflux transporter activities^10,29^.

We set out to compare iBMECs obtained by neuroendothelial differentiation to endothelial cells generated by transition through a mesodermal stage (iECs), referred to as mesoendothelial differentiation^31^. Using a combination of single cell RNA sequencing, bioinformatics and immunofluorescence, we find that iBMECs obtained by neuroendothelial differentiation lack canonical endothelial cell markers (CDH5, PECAM1, VEGFR2, APLNR, eNOS) as well as the ETS transcription factors (ETS1, ETV6, FLI1) crucial for establishing the endothelial identity, when compared to their mesoendothelial counterparts. Despite having some features of endothelial cells (e.g. high TEER), iBMECs do not form lumenized vessels in immunocompromised mice (NOD-scid IL2Rg^null^-SGM3). Moreover, at a molecular level, iBMECs express high levels of EpCAM and exhibit a gene signature typical of neuroectodermal epithelial cells. Analysis by both global and single-cell RNA sequencing combined with immunofluorescence reveals multiple neuroectodermal epithelial related genes^32,33^ such as TTR, TRP3, NKCC and AQP1 present in EpCAM^+^ NE-EpiCs cells.

Finally, NE-EpiCs can be directed towards a BBB-like EC fate by overexpressing three key endothelial ETS transcription factors ETV2, FLI1, and ERG^34^. Thus, similar to hematopoietic stem and progenitor cells^35,36^, directed differentiation of pluripotent stem cells into human BMECs with appropriate neurovascular barrier properties requires introduction of key EC TFs to develop a robust human BBB model *in vitro* that could ultimately be utilized for functional studies and drug discovery.

## Results

### Neuroendothelial differentiation derived iBMECs are not phenotypically endothelial cells

Several groups have developed strategies that simultaneously differentiate human pluripotent stem cells (hPSC) to both neural and EC lineages, rationalizing that this co-differentiation yields hPSC-derived ECs possessing BBB attributes (Fig. 1a, referred to as neuroendothelial iBMECs)^10–22^. To assess the properties of these cells, we compared this protocol to a well-established method in which ECs are differentiated from pluripotent sources through a mesodermal lineage (referred to as mesoendothelial iECs; Fig 1a)^31^, using two IPS cell lines (IMR90-4 and H6) and one ES cell line (H1). The mesoendothelial protocol drives hPSC differentiation towards mesodermal lineages using a chemically defined, serum-free method that enhances vascular differentiation. Heterogenous embryoid body cultures of hPSCs are stimulated with Bone Morphogenetic Protein 4 (BMP-4), Activin A, Fibroblast Growth Factor 2 (FGF-2) and VEGF-A, with the addition of the Transforming Growth Factor beta (TGF-β) inhibitor SB431542^31^ (Fig. 1a). This approach generates iECs that express VE-Cadherin (VECAD; CDH5) and PECAM1 (CD31) but lack EpCAM (CD326) and haematopoietic markers (VECAD^+^PECAM1^+^EpCAM^−^CD45^−^ cells; Fig. 1c, d).

**Figure 1.**
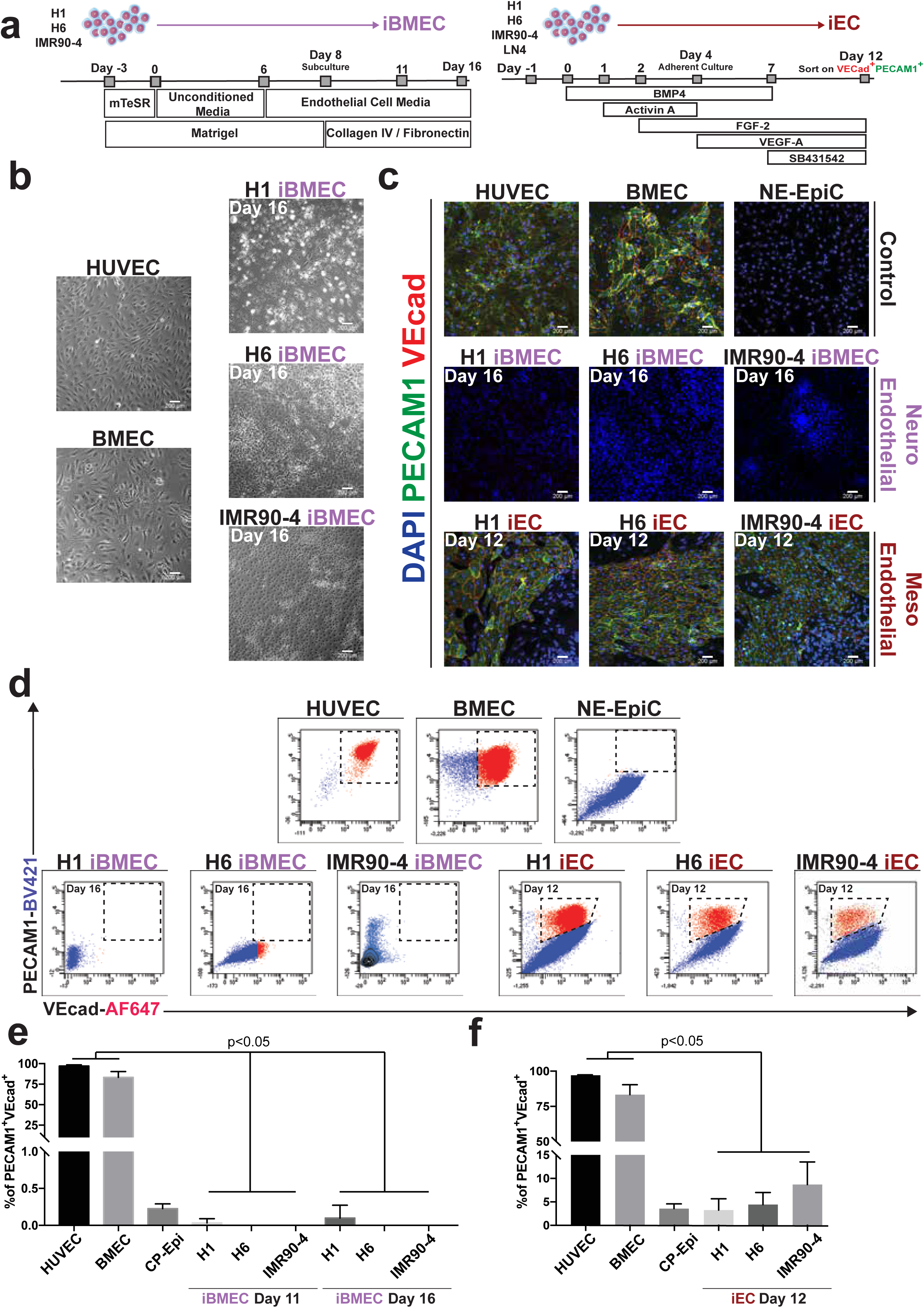
Phenotypic comparison of hPSC-derived cells to *bona fide* endothelium. **a.** Diagrams for differentiation of hPSCs to neuroendothelial iBMECs and mesoendothelial iECs. **b.** Phase contrast microscopy of primary endothelial cells (HUVEC and BMEC) and neuro-endothelial iBMECs derived from H1, H6, and IMR90-4 cell lines at day 11 of differentiation. Representative pictures of n=5 biological replicates, scale bars, 200µm. **c.** Confocal microscopy of HUVECs, BMECs, and NE-EpiCs; H1, H6, and IMR90-4-derived iBMECs at day 16 of differentiation; H1, H6, and IMR90-4 derived iECs at day 12 of differentiation; DAPI (Blue), PECAM1 (Green), and VECAD (Red). Representative pictures of n=5 biological replicates, scale bars, 200µm. **d.** Representative flow cytometry plot assaying VECAD^+^ PECAM1^+^ phenotypically marked cells. n=5 biological replicates. **e.** Quantification of VECAD^+^PECAM1^+^ populations based on FACS analysis.

Phase-contrast microscopy reveals that iBMECs exhibit a characteristic EC morphology (Fig. 1b, Extended Data Fig. 1a; n = 5 biological replicates), however they lack the canonical EC markers VECAD and PECAM1 by day 16 (Fig. 1c, Extended Data Fig. 1b; n = 5 biological replicates). Few iBMECs derived from hPSCs express low levels of VECAD early during differentiation; however, they lose VECAD expression by day 16 (Fig. 1c, d; Extended Data Fig. 1b). Moreover, the VECAD is not homogenously distributed throughout the cell membrane in iBMECs. In contrast, the mesoendothelial-derived iECs, primary Human Umbilical Vein Endothelial Cells (HUVECs) and Brain Microvascular Endothelial Cells (BMECs) express high levels of junctional VECAD and PECAM1 (Fig 1c-d, Extended Data Fig. 1b). Quantification of ECs by fluorescence activated cell sorting (FACS) shows that the neuroendothelial differentiation protocol yields on average 0.03±0.02%, 0.01±0.01%, 0.01±0.01%, VECAD^+^PECAM1^+^ marked cells at day 11 and 0.13±0.07%, 0.02±0.01%, 0.03±0.015% VECAD^+^PECAM1^+^ marked cells at day 16 (for H1, H6 and IMR90-4, respectively; Fig. 1d, e). We tested the in vivo angiogenic potential of iBMECs using matrigel plugs implanted in immunocompromised animals (NSG-SGM3, n=10 animals per cell line tested). Day 16 iBMECs derived from H1, H6 or IMR90-4 fail to establish a vascular network compared to the bona fide endothelial cells (HUVECs) (Extended Data Fig.1c-e). The poor yield of VECAD^+^PECAM1^+^ in iBMECs is not due to the lack of endothelial “competence” from the source cell lines (H1, H6 and IMR90-4), because they produce 3.23±2.41%, 4.43±1.47%, 8.70±2.78% VECAD^+^PECAM1^+^ iECs when differentiated with the mesoendothelial protocol (H1, H6 and IMR90-4, respectively; Fig. 1d, f). Upon purification by FACS on day 12, these iECs can be readily expanded (data not shown). These findings were compared to primary human umbilical vein ECs (HUVECs) and brain microvascular ECs (BMECs), which contain 97.17±0.98% and 83.37±4.028% VECAD^+^PECAM1^+^ cells (Fig. 1d-f). Thus, neuroendothelial differentiation conditions do not generate iBMEC cells with an endothelial phenotype, raising questions regarding their identity.

### The neurorendothelial iBMECs transcriptome is not congruent with an EC signature

We tested the putative iBMEC EC identity using single-cell RNA sequencing (sc-RNAseq). We sequenced and retained after quality control 2812, 2423 and 4980 iBMECs (from H1, H6 and IMR90-4, respectively) plus 4833 iECs (all cell lines included) and 3219 primary ECs (HUVECs/BMECs) (Extended Data Fig. 2b). Single cells transcriptomes were sequenced with an average depth of 7310 nUMI and 2684 genes per cell (Extended Data Fig. 2b). To detect relationships among cells from various differentiation protocols and sources, we visualized all cells with *t*-SNE and grouped them using unbiased hierarchical clustering (Fig. 2a). Because most cells from a similar differentiation protocol cluster together across cell source and biological replicates, this indicates that batch effects are not a source of variance in the dataset (Fig. 2a). To define cell identities, we investigated how the most highly variable genes contribute to cell-type identity, by clustering averaged gene-expression profiles for each cell type using only the top 400 highly variable genes from the dataset (Fig. 2a, cluster a to i’). This shows that manual annotation (Fig. 2a, sample name) of cell types is consistent with unbiased transcriptomic clustering (Fig. 2a, cluster a to i’), suggesting that the effect of cell type, differentiation protocol and cell source on measured gene expression is stronger than batch effects.

**Figure 2.**
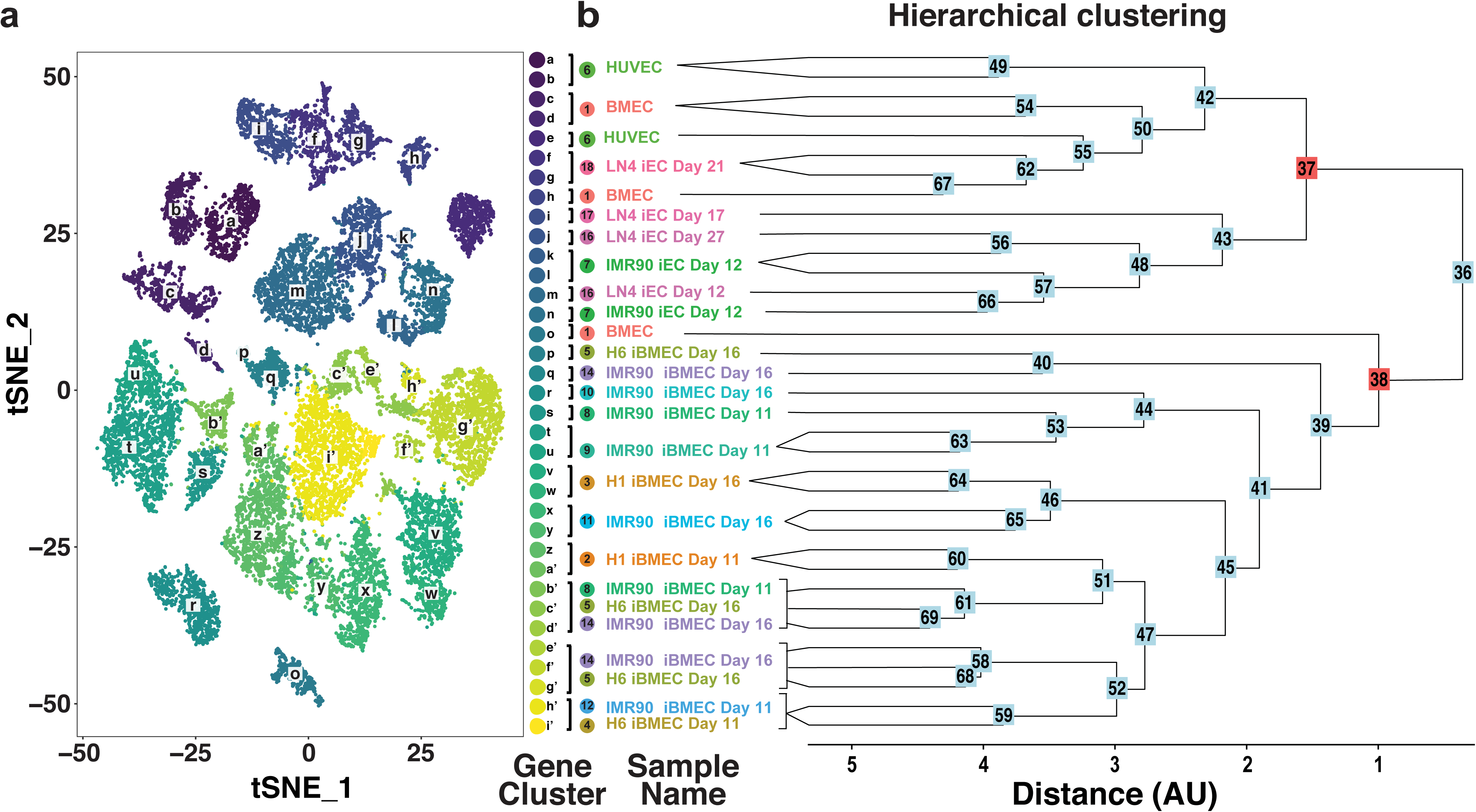
The neuroendothelial iBMECs transcriptome is not endothelial. **a.** scRNA expression profiles displayed on a t-distributed stochastic neighbor embedding (t-SNE) plot illustrates multiple unique clusters (a-i’) of cells separated based on top 400 highest differentially expressed genes. **b.** Dendrogram generated using hierarchical clustering of unique cell clusters represented in Fig. 2a showing a clear distinction between iBMECs and all other endothelial cell clusters.

The resulting dendrogram (Fig. 2b) segregates into two main branches: node #37 and #38 (Fig. 2b). Node #37 accounts for gene clusters a-n and #38 for gene clusters o-i’. Manual annotation revealed that node #37 is comprised of all primary ECs (HUVECs and BMECs) as well as mesoendothelial iEC samples, thereby defining an endothelial core population, while node #38 contains neuroendothelial iBMECs exclusively (Fig. 2a, b). ECs are defined not only by their ability to form blood vessels, but also by expression of unique transcription factors (TFs), signaling pathways and growth (angiocrine) factors^37–40^. Using our unsupervised single cell RNA-seq analysis, we curated a list of essential genes associated with EC identity that are highly expressed by cell clusters a-e from Fig. 2a. This supervised analysis revealed that iBMECs, independently of the cell line and the day of differentiation tested, lack expression of EC transcription factors (ETS1, ETS2, ELK3, ELF1, GATA2, FLI1, SOX17, SOX18) compared to mesoendothelial iECs and *bona fide* adult endothelial cell controls (Fig. 3). iBMECs also lack general EC cell signaling machinery, including VEGFR2 (VEGF pathway), ICAM2, Notch ligands (JAG1, JAG2, NOTCH4, EGFL7) and NOS3 (Fig. 3, Extended data Fig. 4,7). Moreover, iBMECs lack secreted or membrane-bound angiocrine factors inherent to EC identity, such as ANGPT2, APLN, BMP4, BMP6, FGF2, IGFBP4 and IGFBP7 (Fig. 3). CNS ECs are characterized by high levels of WNT/β-catenin signaling which is required to induce and maintain BBB properties^41–44^. We analyzed expression of WNT/β-catenin signaling components in iBMECs and found that they express low levels of key receptors (FZD4, LRP5/6) but lack downstream targets that are indicative of pathway activation (LEF1, NKD1/2, AXIN2, SOX17) (Extended Data Fig. 5). Therefore, the neuroendothelial differentiation protocol yields iBMECs cells that lack both phenotypic and molecular signatures of both general EC lineage and the CNS endothelium.

**Figure 3.**
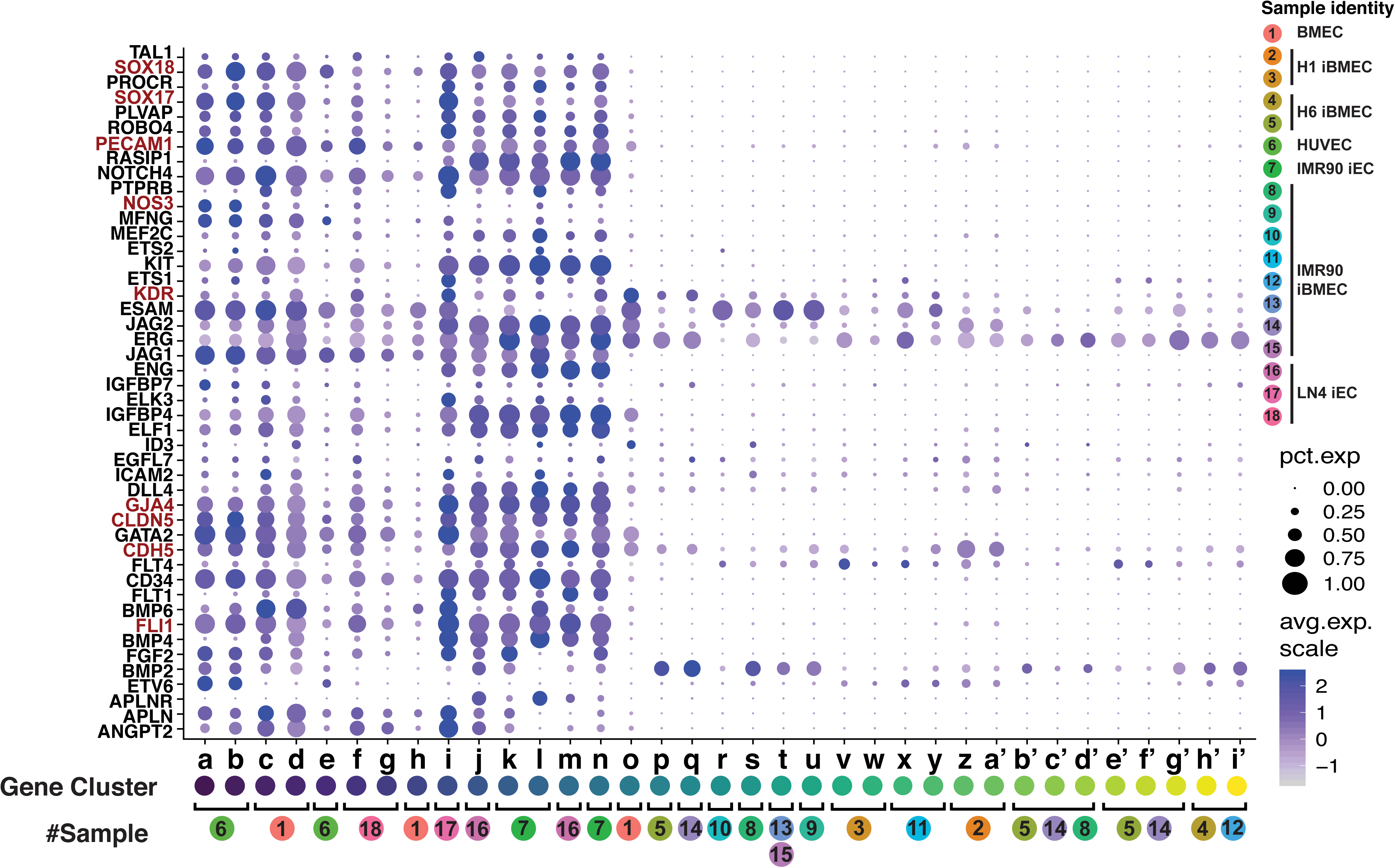
Neuroendothelial iBMECs lack angiocrine factors. Dot plot emphasizing large difference in expression of vascular genes between Endothelial cell population and hPSC-derived iBMECs. For this and all subsequent dot plots, cell clusters and sample numbers are taken from Fig. 2a. Dot size indicates percentage of cells in populations expressing the gene whereas intensity of color dictates expression level.

### iBMECs are EpCAM^+^ cells displaying neuroectodermal epithelial-like properties

Upon further interrogation of our single-cell RNAseq dataset comprising 20,069 cells (Fig. 2a), we identified several genes differentially expressed at high levels between cell populations stemming from nodes 37 and 38 (Fig. 2b). Using these highly differentially expressed genes, we performed a supervised analysis and found that iBMECs express high levels of genes not generally associated with EC in addition to a unique CLAUDIN repertoire. These included EPCAM, KRT7/8/19, SLC12A2, FREM2, and SPP1 (Fig. 4a). Confocal microscopy and FACS analysis both confirm EPCAM expression in a large subset of iBMECs (Fig. 4b). Expression of these genes suggest that iBMECs have an epithelial phenotype in contrast to an endothelial one as previously suggested^10–22^.

**Figure 4.**
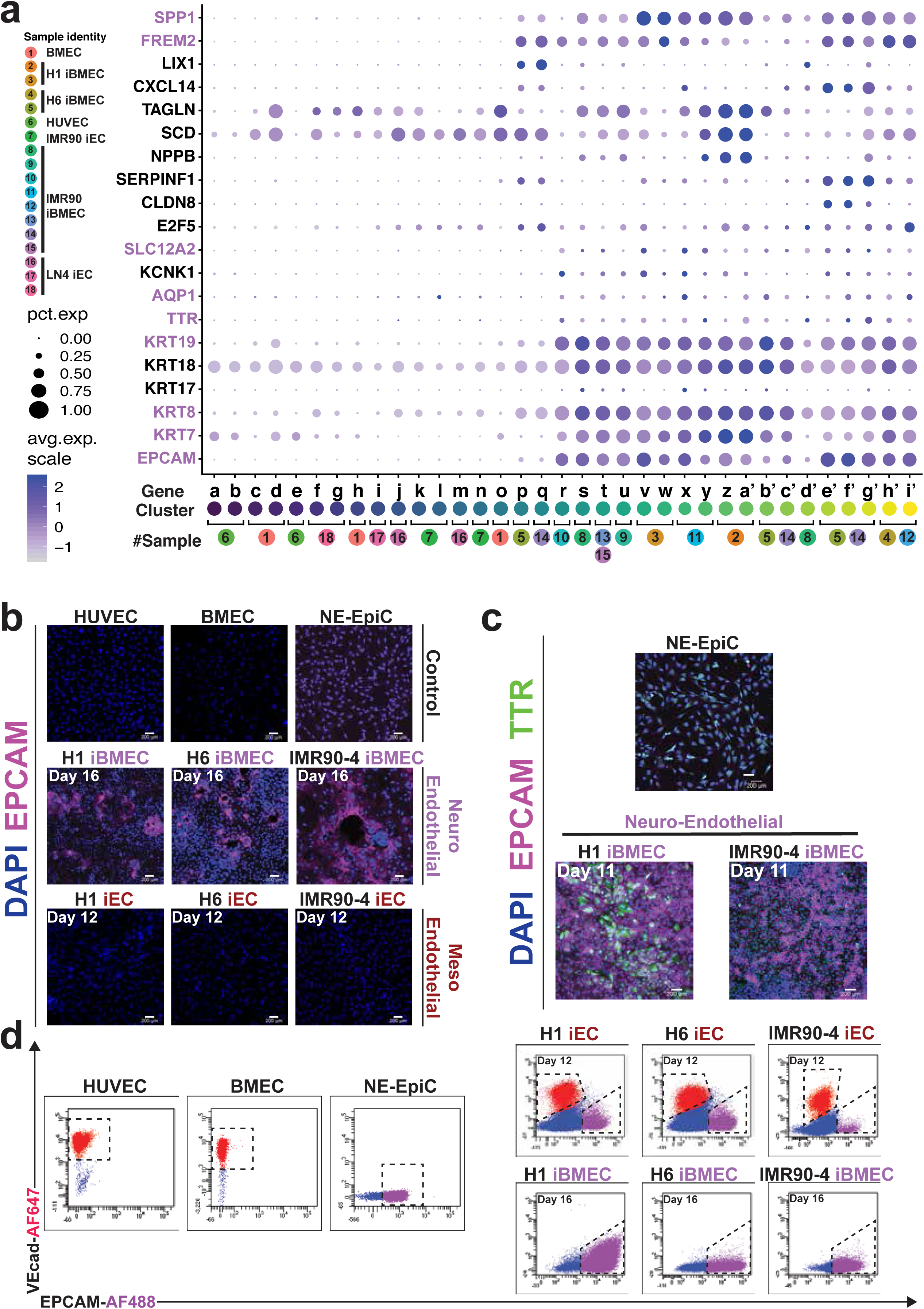
Neuroendothelial iBMECs express genes characteristic of epithelial cells. **a.** Dot plot showing a clear bias in the expression of epithelial-associate genes towards iBMEC samples compared to all endothelial control samples and mesodermal differentiation derived iECs. Highlighted genes correspond to subset of epithelial genes highly expressed in iBMEC cell clusters compared to human ECs. **b.** Confocal microscopy of HUVECs, BMECs, and NE-EpiCs; H1, H6, and IMR90-4 derived iBMECs at day 16 of differentiation; H1, H6, and IMR90-4 derived iECs at day 12 of differentiation. DAPI (Blue) and EpCAM (Purple). Representative pictures of n=5 biological replicates, scale bars, 200µm. **c.** Confocal microscopy showing NE-EpiCs and H1 and IMR90-4 derived iBMECs. DAPI (Blue), EpCAM (Purple), TTR (Green). n=5 biological replications, scale bars, 200 µm. **d.** Representative flow cytometry plot assaying VECAD^+^EpCAM^+^ phenotypically marked cells. n=5 biological replicates.

Neurovascular ECs form the BBB through expression of several tight junction proteins (CLAUDIN-5, OCCLUDIN, ZO-1/2)^3^. To assess the BBB-like signature of iBMECs, we mined our dataset using the KEGG tight junction pathway (map04530). Our analysis reveals only minor gene expression differences among iBMECs, mesoendothelial iECs and EC controls (HUVEC, BMEC) with the exception of the CLAUDIN family repertoire (Extended Data Fig. 6, 7). All iECs and primary EC controls express CLAUDIN-5 and -11, while iBMECs express a distinct repertoire (CLAUDIN-4, 6 and 7), which is characteristic of epithelial cells^45^ (Extended Data Fig. 6). However, in contrast to epithelial cells or BBB-forming ECs, iBMECs express low levels OCCLUDIN, JAM2 and JAM3 at the RNA level (Extended Data Fig. 6). We performed immunofluorescence for CLAUDIN-5, OCCLUDIN and ZO-1 proteins in these cells using the published antibodies^10,29^. We found that antibodies for CLAUDIN-5 and OCCLUDIN recognize proteins at cell-cell junctions that co-localize with ZO-1 (Extended Data Fig. 8a-f). We then measured TEER over time and found that these cells form high TEER (Extended Data Fig. 8g,h) consistent with previous studies^10,29^. However, given that CLAUDIN-5 and OCCLUDIN are not present at the RNA level in these cells (Extended Data Fig. 6), we conclude that these antibodies may cross-react with epithelial CLAUDIN proteins. Therefore, the neuroendothelial differentiation protocol generates iBMEC cells that do not exhibit the prototypical tight junction protein repertoire normally found in either peripheral or CNS ECs.

Within the CNS environment, the choroid plexus contains epithelial cells that form tight junctions and prevent transcytosis of unwanted substances from the blood into the cerebrospinal fluid (CSF), to form the blood-CSF barrier^20,46–48^. We investigated whether iBMECs resemble these neuroectodermal epithelial cells (NE-EpiCs). Single-cell RNAseq analysis shows that iBMECs express some of the genes associated with the ependymal cell lineage of the choroid plexus such as EPCAM, KRT7/8/18/19, SLC12A2, FREM2 and SPP1 (Fig. 4a,b,d). Additionally, iBMECs express low levels of some choroid plexus epithelial genes such as TTR, AQP1, KRT17, or CLDN1 (Fig. 4a, Extended data Fig. 6). Significantly, none of these epithelial genes were present in either iECs generated from the same hPSC cell lines or control endothelium (Fig. 4a). We also validated the expression of transthyretin (TTR), a terminal differentiation marker of choroid plexus epithelial cells, by immunofluorescence. We found that some EpCAM^+^ cells express TTR, suggesting an ependymal/choroid plexus-like identity (Fig. 4c).

### The neuroendothelial differentiation protocol reproducibly generates non-neural epithelial cells lacking endothelial identity

To substantiate the validity of our analysis, global gene expression profiles obtained by bulk RNA sequencing of previously published iBMECs^18^ were compared to our iBMECs and *bona fide* ECs (HUVECs and BMECs plus previously published iBMECs ^18,34^; Fig. 5a). Hierarchical clustering based on the top 500 highly variable genes across the dataset reveals clear segregation between iBMECs and *bona fide* ECs (Fig. 5a). Notably, a previously published primary BMEC control (Qian *et al.*^18^) clusters with our endothelial cell control, strongly suggesting that segregation between iBMECs and ECs is not due to either library preparation or sequencing bias. We then supervised this clustering using the genes that define our epithelial/endothelial groups from single-cell RNA-seq analysis. iBMECs derived from neuroendothelial co-differentiation show a distinct separation from all adult ECs, regardless of the source of the laboratory that generated them^10,18,29^ (Fig. 5b). Previously published lines^10,18,29^ as well as iBMECs generated by a chemically defined differentiation protocol^18^ lack key endothelial surface markers (CDH5, PECAM1, CLDN5, ICAM2, EGFL7), critical angiocrine factors (NOS3, BMP6, BMP2, KDR) as well as ETS transcription factors (FLI1, ERG, SOX17, TAL1) that are fundamental for endothelial identity (Fig. 5b). Taken together, these data strongly suggest that the custom two-dimensional hPSC differentiation strategy promoting neural and “endothelial” co-differentiation described in^10,18,29^ reproducibly generates pure populations of neuroectodermal epithelial cells lacking an endothelial identity.

**Figure 5.**
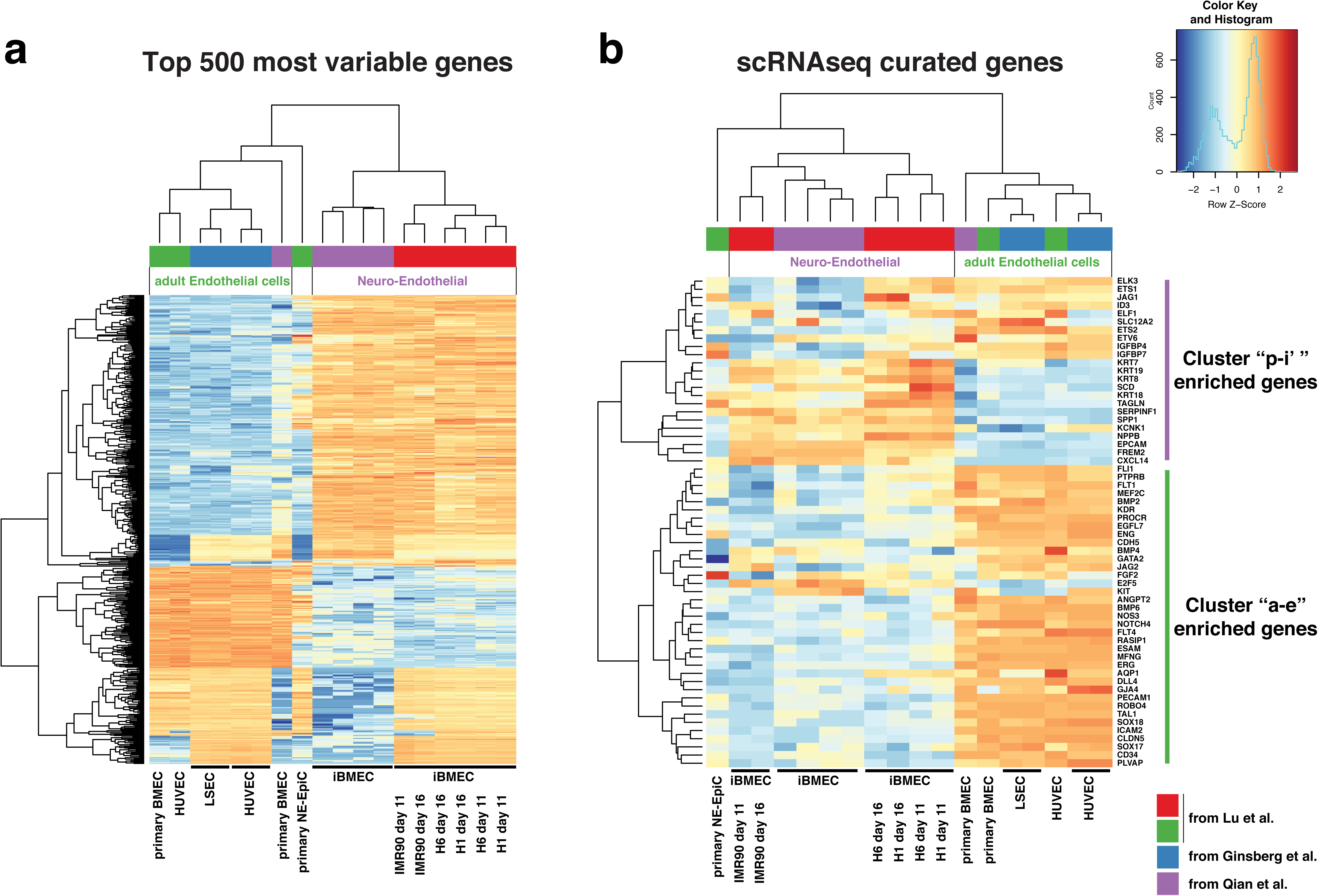
Global RNA sequencing heat maps. **a.** Heat map of top 500 most variable genes between all cell samples both previously published or acquired in house. **b.** Heat map of endothelial and epithelial genes derived from the single cell dataset expressed by all cell samples both previously published or acquired in house.

### ETS transcription factors drive neuroectodermal epithelial cells towards an endothelial fate

We previously showed that lineage-committed epithelial cells (EpCAM^+^Tra1-81^−^c-Kit^−^) from amniotic fluid are amenable to transcription factor-mediated reprogramming into vascular ECs^34^. Transient ETV2 along with constitutive FLI1 and ERG1 expression not only stably induces expression of EC genes in committed epithelial cells, but also suppresses expression of non-vascular genes^34^. We therefore transduced IMR90-4 iPS cells with the ETS transcription factors ETV2, ERG and FLI1 (EEF) at day 6 of the neuroendothelial co-differentiation protocol when epithelial fate is induced (Fig. 6a). We allowed these TFs to be expressed for the duration of differentiation (Fig. 6a). Under these conditions, EEF-iBMECs express both PECAM1 and VECAD by day 11 (Fig. 6b). These cells can be sorted and, with the addition of a TGF-β signaling inhibitor (SB431542), expanded as stable, phenotypically marked ECs defined by VECAD^+^PECAM1^+^EpCAM^−^ expression by day 27 (Fig. 6b, lower panel). Moreover, EEF-enforced expression followed by TGF-β inhibition restores VECAD and PECAM1 proper localization to cell junctions (Extended data Fig. 3b). We inspected the quality of our reprogramming strategy using single-cell RNA sequencing. Compared to untransduced controls, day 11 EEF-iBMECs show tight association with iECs, HUVECs and primary BMECs (Fig. 6c-d). A significant number of vascular genes (VECAD, PECAM1, CLND5 and SOX17) are upregulated in EEF-iBMECs compared to untransduced control iBMECs (Fig. 6c). Expression levels of these EC genes are comparable to those from iECs, HUVEC and primary BMECs cultured *in vitro*, supporting the notion that EEF-iBMECs have an endothelial identity. Moreover, angiocrine factors that regulate EC function^40^ such as BMPs and Notch ligands are all expressed in EEF-iBMECs (Fig. 6c). In conclusion, genome-wide, single-cell RNA sequencing analyses demonstrate that forced expression of ETV2, FLI1 and ERG1 in conjunction with TGF-β inhibition reprograms neuroectodermal epithelial cells into endothelial cells while silencing non-vascular genes, strengthening our assertion that the chemically defined neuroendothelial differentiation protocol^10,18,29^ fails to produce CNS-specific endothelium with BBB properties.

**Figure 6.**
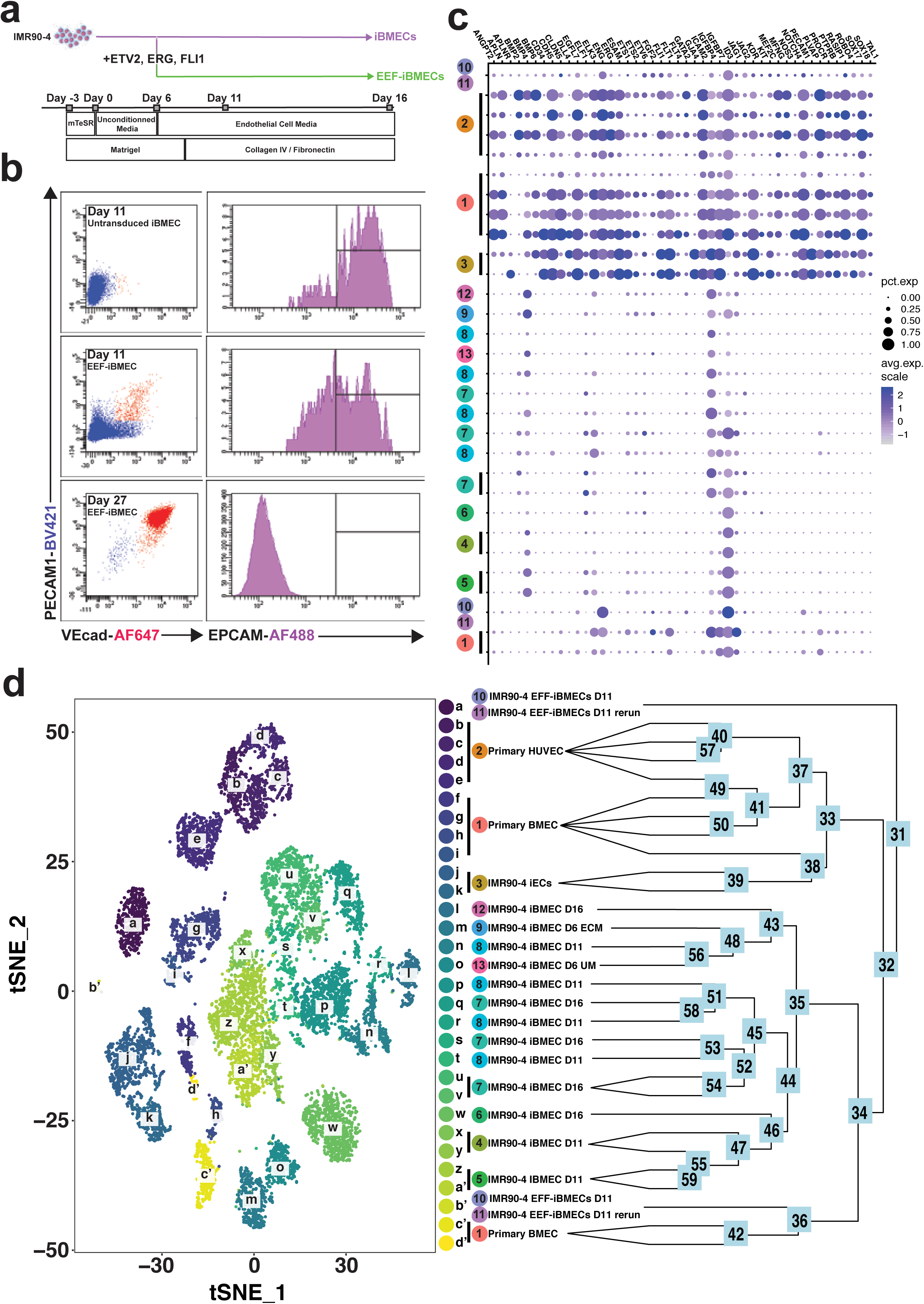
Generating phenotypic ECs from iBMECs using ETV2, ERG and FLI1. **a.** Schematic for the generation of EEF-iBMECs highlighting the induction of ETV2, ERG, FLI1 at day 6 of the neuro-endothelial codifferentiation. **b.** Representative flow cytometry plot assaying VECAD^+^PECAM1^+^ phenotypically marked cells as well as EpCAM expression. Top panels; Untransduced control iBMEC sample, Middle panels; iBMEC transduced with EEF factors at day 6 of neuro-endothelial codifferentiation, Bottom panels; EEF-iBMECs at day 27 (post day 11 sorting). n=5 biological replicates. **c.** Dot plot showing expression levels and prevalence of vascular genes between primary ECs (HUVECs, BMECs), iECs, and IMR90-4 derived cells (iBMECs, EEF-iBMECs). **d.** Left; scRNA expression profiles displayed on a t-distributed stochastic neighbor embedding (t-SNE) plot illustrates multiple unique clusters (a-d’) of cells separated based on top 500 highest differentially expressed genes. Right; Dendrogram generated using hierarchical clustering of unique cell clusters.

## Discussion

Contrary to several prior reports^10–22^, here we demonstrate that iBMECs generated by neuroendothelial co-differentiation from H1, H6 and IMR90-4 hPSCs are not proper BBB-forming ECs, but rather EPCAM^+^PECAM1^−^VECAD^−^ neuroectodermal epithelial cells with similarities to the choroid plexus cells, rendering their use as an *in vitro* model of the BBB questionable.

Our analysis also illustrates that iBMECs are fundamentally different from iECs arising from the same cell lines derived through mesodermal differentiation, as well as primary HUVEC and BMECs. iECs derived from a mesodermal differentiation protocol are EPCAM^−^PECAM1^+^VECAD^+^ cells that can expand *in vitro* and express *bona fide* ECs characteristics. Therefore, there is no inherent restriction of human iPSC or ESC cells to acquire EC properties. An unbiased, graph-based clustering of our single-cell RNAseq data shows that mesodermal-derived iECs closely resemble primary human ECs while neuroendothelial iBMECs do not, further underscoring the vast transcriptional and phenotypic differences that exist between iBMECs and CNS ECs. Upon manual annotation of these unbiased cell clusters with their corresponding sample identities, we conclude that batch effect and starting cell line are not the main source of variance in these results. Moreover, bulk RNA sequencing, which has a greater depth of sequencing, confirms the identity of the iBMECs obtained from our single cell RNA sequencing data. In an unbiased analysis using bulk RNA seq data, our iBMECs were found to have nearly identical global gene expression profiles as cells generated from the neuro-endothelial co-differentiation in several previous studies^10,18,29^. These findings substantiate our conclusions that the published neuroendothelial protocol is working in our hands. The same analysis also shows a primary BMEC control from the same previously published dataset which closely clusters with our primary EC controls, allowing us to conclude that sequencing bias and batch effect do not account for any major differences in the assigned cell identities. Despite the presence of “BBB-like” properties such as the presence of tight junction proteins, ability to form a high TEER and expression of several transporters due to their epithelial nature, iBMECs cannot be used as an *in vitro* model of the human BBB for future studies.

We have not shown, however, that iBMECs are definitive hPSC-derived neuroectodermal epithelial-like cells (NE-EpiCs), since they harbor many, but not all attributes of an epithelial lineage present in the CNS. We demonstrate that constitutive overexpression of ETS factors (FLI1, ERG, ETV2) using lentiviral vectors during the specification phase, reprograms NE-EpiCs towards an EC identity as demonstrated by an EPCAM^−^PECAM1^+^VECAD^+^ phenotype. Although these cells may not represent definitive BBB-forming CNS ECs, the EEF-iBMECs have a more valid EC identity than the iBMECs currently in use as an *in vitro* BBB model. These reprogrammed cells have both the capacity to be passaged and expand *in vitro*, while retaining their PECAM1^+^VECAD^+^ EC phenotype. We believe that this is a crucial step towards *de novo* generation of true BBB-forming CNS ECs for suitable *in vitro* modeling of physiological and pharmaceutical studies which remains a major issue with the current iBMECs. Therefore, while neuroendothelial differentiation of human iPCS to BMECs is rare, overexpression of ETS transcription factors may promote conversion into vascular cells that are capable to form barrier properties.

## Methods

### Maintenance of hPSCs and neuroendothelial co-differentiation to iBMECs

These protocols were precisely followed as described previously^10,28,29^. All reagents were procured based on these protocols, and each step and duration for differentiation was followed rigorously. IMR90-4 and H6 iPSCs and H1 hESCs were maintained between passages 20–42 on Matrigel (BD Biosciences, 354277) in mTeSR1 or TeSR-E8 medium (STEMCELL Technologies, 85850, 05990). For differentiation, cells were passaged onto Matrigel in mTeSR1 or StemFlex (Gibco, A3349401) medium and allowed to expand for 3 days. Cultures were then switched to unconditioned medium (UM) lacking bFGF for 6 days. EC medium consisting of human endothelial serum-free medium (hESFM; Gibco, 11111044) supplemented with 20 ng/mL bFGF (Peprotech, 100-18B), 1% platelet-poor plasma-derived bovine serum (Alfa Aesar, J64483AE), and 10 μM all-trans retinoic acid (Sigma, R2625) was then added for an additional 2 days. Cells were then dissociated with Accutase (Invitrogen, 00-4555-56) and plated onto 6-well polystyrene plates or 1.12 cm^2^ Transwell-Clear permeable inserts (0.4 mm pore size) in EC media. Culture plates were coated with a mixture of Collagen IV (400 mg/mL; Sigma, C6745) and Fibronectin (100 mg/ mL; Sigma, F1141) in H_2_O for at least 30 min at 37°C, whereas inserts were incubated for a minimum of 4 h at 37°C. The resulting purified hPSC-derived iBMECs were then grown in EC medium for 24 h, after which RA was removed and cells were cultured until the indicated experimental timepoints.

### PSC mesoendothelial differentiation to iECs

hPSC were maintained exactly as described above. Embryoid bodies were generated and cultured in base hPSC medium consisting of StemPro-34 (Gibco, 10639011), nonessential amino acids (Gibco, 11140050), glutamine (Gibco, 35050061), penicillin/streptomycin/amphotericin B (Gibco, 15240096), β-mercaptoethanol (MilliporeSigma, M3148) for 24 hours before the start of differentiation and the protocol was carried out as previously described^31^. Medium is changed every 2 days for the duration of the differentiation. Addition of 20 ng/ml BMP-4 (R&D Systems, 314-BP-05/C) on day 0 (removed on day 7); on day 1, medium is supplemented with 10 ng/ml activinA (STEMCELL technologies, 78001; removed at day 4); on day 2, medium is further supplemented with 8 ng/ml FGF-2 (Peprotech, 100-18B; remained for the duration of culture); on day 4, embryoid bodies are transferred to adherent conditions on Matrigel-coated plates and medium was supplemented with 25 ng/ml VEGF-A (Peprotech, 100-20; remained for the duration of culture) to specify vascular progenitors; on day 7, SB431542 (R&D Systems, 1614; TGFβ signaling inhibitor) is added at 10 μM concentration and remained for indicated duration to expand terminally differentiated endothelial cells (iECs)

### Directing hPSC neuroendothelial co-differentiation towards EEF-iBMECs

IMR90-4, H6, and H1 hPSCs were differentiated according to the Neuro-Endothelial co-differentiation protocol as described previously^34^. Following the 6 days of differentiation in UM, cells were transduced via lentiviral vectors for ETS transcription factors FLI1, ERG, ETV2 and cultured in EC medium, exactly as mentioned above. No further changes were made to the protocol because the cells were terminally cultured.

### Single-cell RNAseq digital droplet sequencing (ddSeq)

A single-cell suspension was loaded into the Bio-Rad ddSEQ Single-Cell Isolator on which cells were isolated, lysed and barcoded in droplets. Droplets were then disrupted and cDNA was pooled for second strand synthesis. Libraries were generated with direct tagmentation followed by 3’ enrichment and sample indexing using Illumina Nextera library prep kit. Pooled libraries were sequenced on the Illumina NextSeq500 sequencer. Sequencing data were primarily analyzed using the SureCell RNA Single-Cell App in Illumina BaseSpace Sequence Hub.

All single cell analyses were performed using the Seurat package in R (version 2.3.4). UMI counts files that had been knee-filtered were downloaded from the Illumina BaseSpace Sequence Hub. After initial quality control, cells that were included in the analysis were required to have a minimum 900 genes expressed and a maximum 5500 genes expressed, in addition to a minimum 1100 UMIs. This resulted in a total 20,069 cells passing quality filters across the 18 samples as seen in Extended Data Figure 2b. Following best practices in the package suggestions UMI counts were log-normalized and after the most highly variable genes selected the data matrices were scaled using a linear model with variation arising from UMI counts mitigated for. Principal component analysis was subsequently performed on this matrix and after reviewing principal component heatmaps and jackstraw plots t-distributed stochastic neighbor embedding analysis (tSNE) visualization were performed on the top 40 components and clustering resolution was set at 1.0 for visualizations. Differential gene expression for gene marker discovery across the clusters were performed using the Wilcoxon rank sum test as used in the Seurat package.

### Immunofluorescence

#### Fixed Samples

Cells were fixed with 4% paraformaldehyde for 15 minutes prior to staining. To prevent non-specific binding of the primary antibody, samples were blocked with 5% horse serum in PBS solution for 60 minutes. Cells were washed 3 times in PBS and dilute primary antibody in 5% horse serum was added according to manufacturer’s recommended concentration and left overnight at 4°C. If the protein was intracellular, permeabilization using 0.2% Triton X-100 was necessary during block and staining with primary antibody. After staining with primary antibody, cells were incubated with secondary antibody in 5% horse serum for 1 hour at room temperature on a shaker. Following a thorough wash to remove unbound secondary antibody, DAPI was used at 1 µg/ml to stain nuclei.

#### Live Samples

Human IgG antibody was used at 1:50 in PBS with 0.5% BSA and 2□mM EDTA to block F_c_ receptors prior to staining. The blocked cells were stained for 30 minutes at 4°C with fluorochrome-conjugated antibodies according to the manufacturer’s recommendations. Stained cells were washed in PBS to prevent saturation of the fluorescent dye during imaging. All imaging was performed using a Zeiss 710 META confocal microscope.

### Flow cytometry

Before staining, F_c_ receptors were blocked with human IgG antibody at 1:50 (Biolegend) in PBS (pH 7.2) containing 0.5% BSA (Fraction V) and 2□mM EDTA for 10□min at 4□°C. For cell surface staining, samples were stained for 30□min at 4□°C with fluorochrome-conjugated antibodies according to the manufacturer’s recommendation. Stained cells were washed in blocking buffer and fixed in 1% PFA in PBS (pH 7.2) with 2□mM EDTA for flow analysis or resuspended in PBS (pH 7.2) with 2□mM EDTA and 1□μg/ml DAPI (Biolegend) for sorting. Samples were analysed using a FACS ARIA II SORP (BD Biosciences) Data were collected and analysed using FACs DIVA 8.0.1 software (BD Biosciences). All gating was determined using unstained controls and fluorescence minus one strategies.

### Bulk RNA sequencing

Total RNA, greater than 100ng, from cultured cells was isolated in TRIzol L and purified using Qiagen RNeasy Mini Kit per manufacturer’s protocols. Agilent Technologies 2100 Bioanalyzer was used to assess the RNA quality. RNA libraries were prepared and multiplexed using Illumina TruSeq RNA Library Preparation Kit v2 (non-stranded and poly-A selection) and 10 nM of cDNA was used as the input for high-throughput sequencing via Illumina’s HiSeq 2500 platform, producing 50 bp paired end reads. Paired end sequencing reads were then quality-checked and trimmed where appropriate using Trimmomatic 0.36^49^ and the resultant filtered reads reads were mapped to the human reference genome GRCh38 using STAR aligner^50^. Aligned bam files were subsequently sorted and raw transcript counts were quantified using HTSeq^51^ to produce counts matrices based off the human reference transcriptome for GRCh38. Previously published RNAseq data for hPSC derived iBMECs and primary BMECs were downloaded from the Gene Expression Omnibus (GEO, GSE97575)^18^ along with previously published data for HUVECs and LSECs (GEO, GSE40291)^52^ and were processed in the same way as for the bulk RNAseq libraries prepared for the experiment. Raw counts were processed in R package edgeR^53^ and a Log2 counts per million matrix was calculated after filtering and quality control. A heatmap displaying the 500 most variable genes was subsequently formulated with the distance between both rows and columns being calculated via Euclidean distance and with dendrograms computed and reordered based on row and column means.

### In vivo tubulogenesis assay

H1, H6, and IMR90-4 derived neuroendothelial iBMECs at day 16 of differentiation and primary HUVECs were resuspended in Matrigel (Corning, 254277) at 500,000cells/300μL and injected subcutaneously into immunocompromised mice (NSG-SGM3; Jax, 013062). Mice were also injected with an empty Matrigel control. After 5 days, mice were sacrificed following institutional guidelines and solidified Matrigel plugs were excised and fixed using 4% paraformaldehyde (Alfa Aesar, AA43368-9M). Plugs were cryosectioned, stained using CD144 (BV9, Biolegend), Agglutinin 1 (UEA1, Vector Labs), and EPCAM (9C4, Biolegend). Slides were fixed using Fluoroshield with DAPI (Sigma-Aldrich, F6057) and imaging was performed using a Zeiss 710 META confocal microscope. Quantification analysis of images were conducted using Angiotool^54^.

### Virus production

Human ETV2, FLI1, and ERG lentiviral particles were generated and tittered as described in Ginsberg *et al.*, with the following modifications. 293T Lenti-X cells (passage 8–10; subconfluent) were used to produce lentiviral particles using Lenti-X packaging single-shot system, following the manufacturer’s protocol. Filter through a 0.45μm PES filter attached to a 30ml syringe and use Lenti-358 X concentrator following manufacturer’s protocol. Resuspend the concentrate from a 100mm plate of 293T cells in 100 μl of lentivirus storage buffer and store at −80°C for up to 1 year.

### Trans-Endothelial Electrical Resistance Measurement

Cells were plated on poly-D-lysine and Collagen IV-coated gold electroarray 96-well plates (Applied Biophysics) and grown in endothelial cell media supplemented with growth factors and 10% FBS media (Cell Biologics) to maximum resistance. Cells were switched to 1% FBS and the resistance recorded every 30 min for 48 hr using the ECIS Z-θ system (Applied Biophysics). Resistance curves were generated using GraphPad Prism software. Areas under the curve were quantified for the final 48 hr after cells reached maximum resistance using GraphPad Prism software.

### Animal experiments

12-15 weeks old immunodeficient mice, specifically NSG-SGM3, Jax, 013062, were used in this study. All animal experiments were performed under the approval of Weill Cornell Medicine Institutional Animal Care and Use Committee (#2009-0061).

## Author contributions

TL, DA, SR and RL designed and conducted various aspects of the experiments wrote and edited the manuscript. DR interpreted the single cell and bulk RNA seq. AS, AnS, TM, DHN, JX provided technical assistance. KS, HAF, ZR helped to formulate the hypothesis, interpret the results, and edited the manuscript. SR, DA & RL directed and supervised all aspects of the study.

## Acknowledgements

We thank Raul Chavez for the technical assistance and the staff in the Genetic Resources Core Facility in Weill Cornell Medicine. We thank Tyler Cutforth at Columbia University Irving Medical Center for his editorial input in the manuscript. D.A. is supported by the National Institutes of Health (#R01MH112849 #R01NS107344), the Leducq Foundation #15CVD-02 and unrestricted gifts from John Castle, Newport Equity LLC and PANDAS Network to the Department of Neurology, Division of Cerebrovascular Diseases and Stroke at CUIMC (D.A.). H.A.F is supported by the National Institute of Health (NIH DP1CA228040), AnS is supported by funding from the Henry and Marylin Taub Foundation. T.L., K.S., S.R. and R.L. are supported by the Ansary Stem Cell Institute, Starr Foundation TRI-Institution Stem Cell Initiative grants (Stem Cell Derivation Core, #2013-032, #2014-023, #2016-013), Qatar National Priorities Research Program (NPRP 8-1898-3-392), the Empire State Stem Cell Board and New York State Department of Health grants (NYSTEM: C026438, C026878, C029156) and by grants from the NIH R01 (DK095039, HL119872, HL115128, RC2DK114777, HL128158, HL139056-01A1, U01AI138329).

---

**Extended Data Figure 1.**
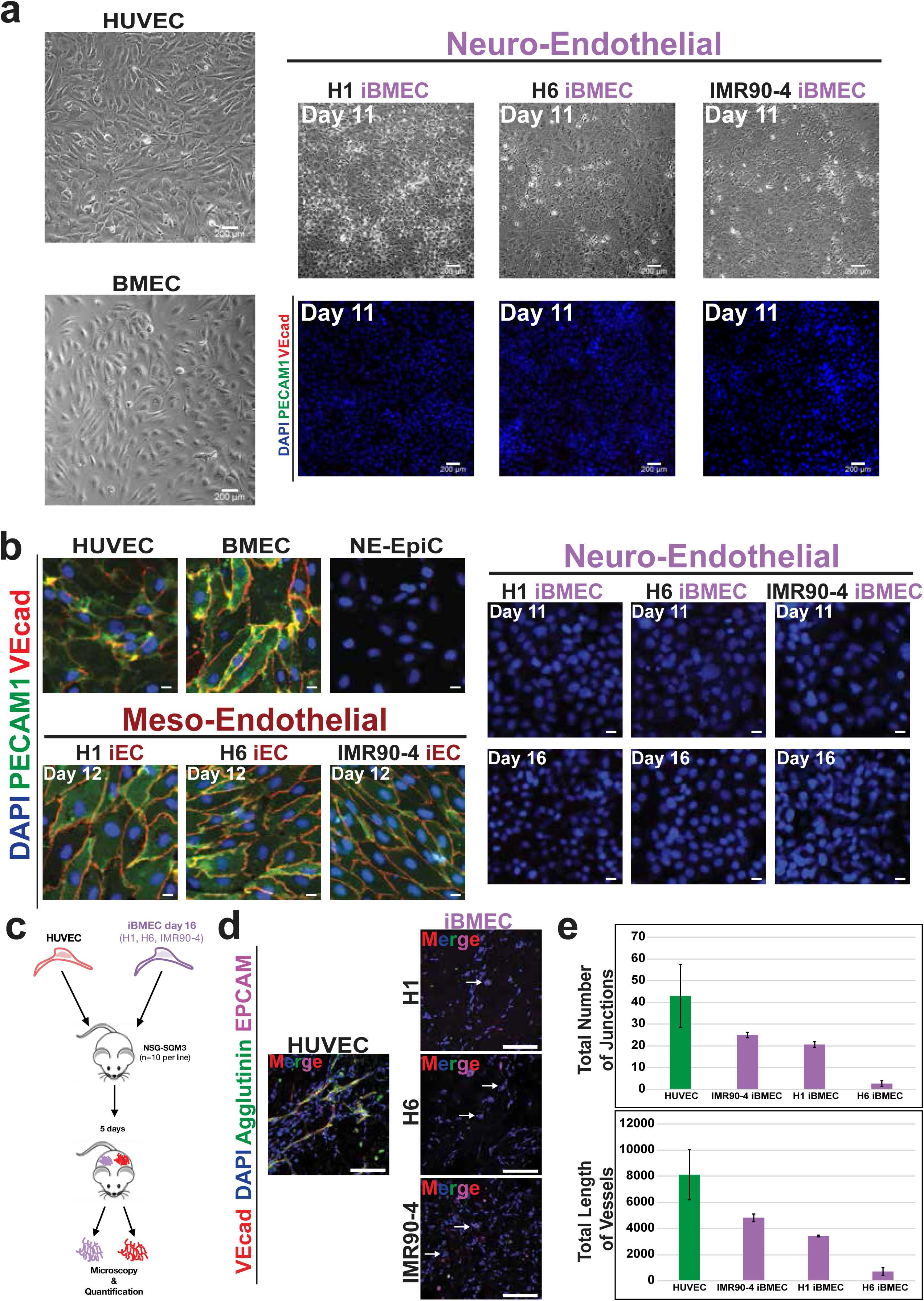
Phenotypic and functional comparative assay of iBMECs. **a.** Phase contrast microscopy of primary endothelial cells (HUVEC and BMEC) and neuroendothelial iBMECs derived from H1, H6, and IMR90-4 cell lines at day 11 of differentiation. Confocal microscopy of H1, H6, and IMR90-4 derived iBMECs at day 11 of differentiation; DAPI (Blue), PECAM1 (Green), VECAD (Red). n=5 biological replicates, scale bars, 200µm. **b.** Confocal microscopy of H1, H6, and IMR90-4 derived iBMECs at day 11 and day 16 of differentiation; H1, H6, and IMR90-4 derived iECs at day 12 of differentiation; HUVECs, BMECs, and NE-EpiCs. DAPI (Blue), PECAM1 (Green), and VECAD (Red). Representative pictures of n=5 biological replicates, scale bars, 50µm. **c.** Schematic of in vivo vessel formation assay in NSG-SGM3 mice. 500,000 iBMEC or HUVEC in Matrigel were injected subcutaneously into 10 mice per hPSC cell line and excised 5 days later for microscopy analysis. **d.** *In vivo* tubulogenesis assay of HUVEC and H1, H6, and IMR90-4 derived iBMECs at day 16 of differentiation in Matrigel plugs. DAPI (Blue), EpCAM (Purple), Agglutinin (Green), and VECAD (Red). White arrows in iBMEC panels identifying groups of iBMECs forming EpCAM^+^ cell clusters. **e.** Angiotool quantifications of *in vivo* tubulogenesis assay, Top; total number of vessel junctions in each sample, Bottom; total length of all vessels in each sample. n=3 biological replicates, 10 mice per cell line tested.

**Extended Data Figure 2.**
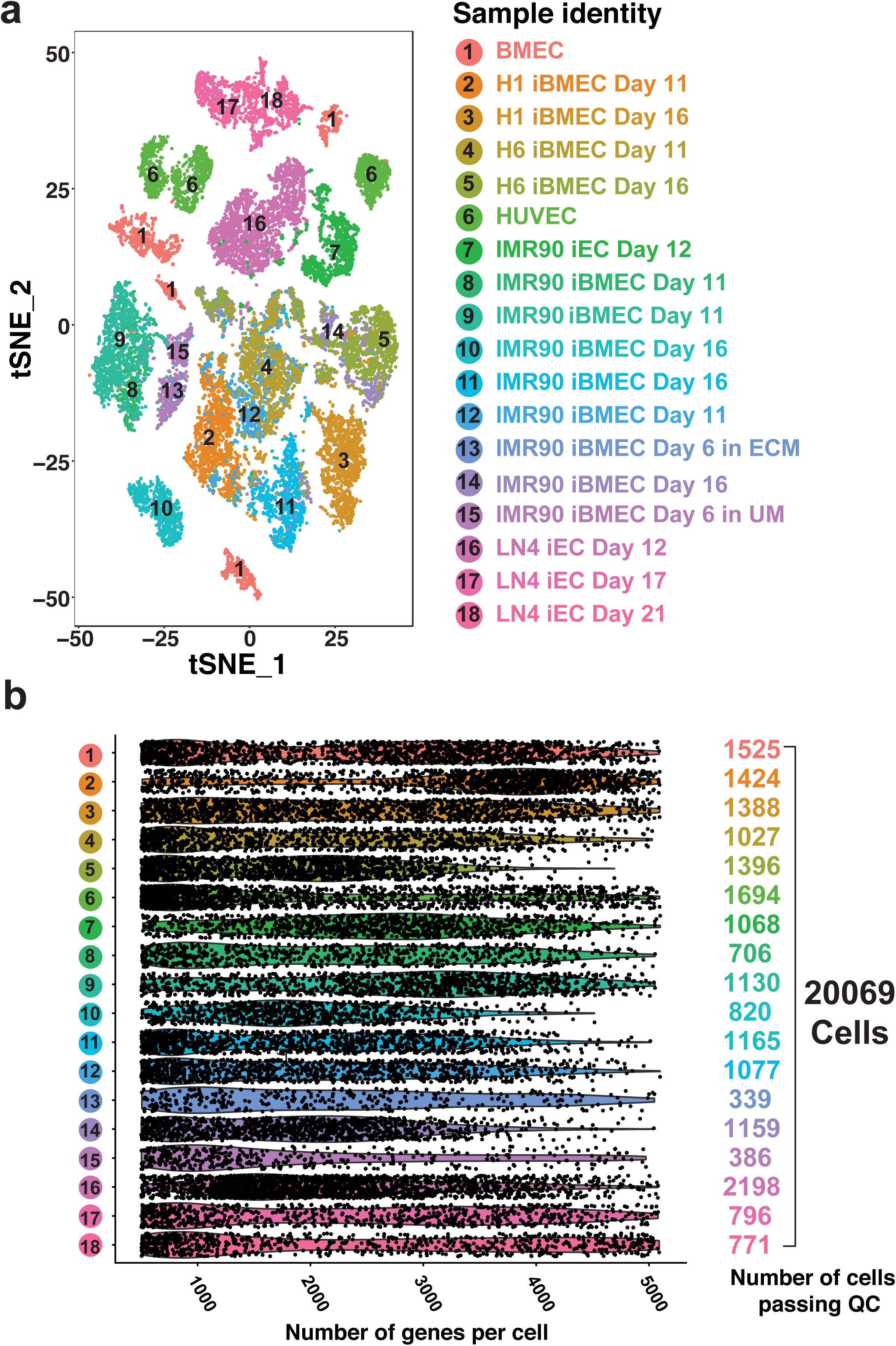
Single-cell RNA sequencing samples and quality control. **a.** scRNA expression profiles displayed on a t-distributed stochastic neighbor embedding (t-SNE) plot illustrates 18 distinct cell samples used in primary analysis of endothelial cells (HUVEC and BMEC), meso-endothelial iECs at day 12, and neuro-endothelial iBMECs at days 11 and 16. **b.** Violin plot describing total number of cells per sample that passed quality control and therefore were used in the study.

**Extended Data Figure 3.**
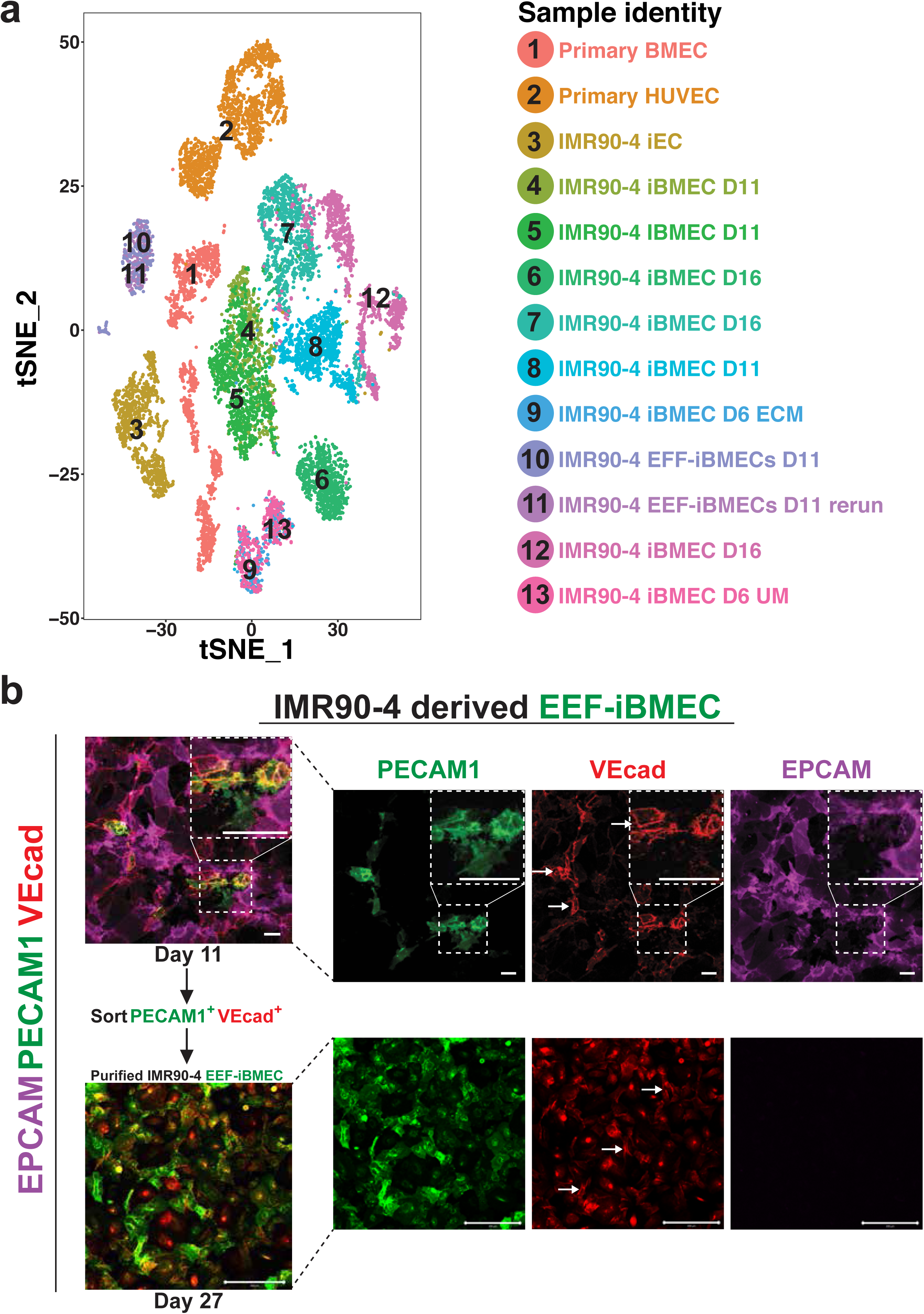
Relative identity of IMR90-4-derived EEF-iBMECs. **a.** scRNA expression profiles displayed on a t-distributed stochastic neighbor embedding (t-SNE) plot illustrates endothelial cells (HUVEC and BMEC), IMR90-4 derived iBMECs at days 11 and 16, and IMR90-4 derived EEF-iBMECs at day 11. **b.** Confocal microscopy of EEF-iBMECs at days 11 and 27 (post day 11 FACS) showing EpCAM (purple), PECAM1 (Green), and VECAD (red). White arrows showing regions of junctional VECAD being restored in EEF-iBMECs. n=5 biological replicates, scale bars, 200µm.

**Extended Data Figure 4.**
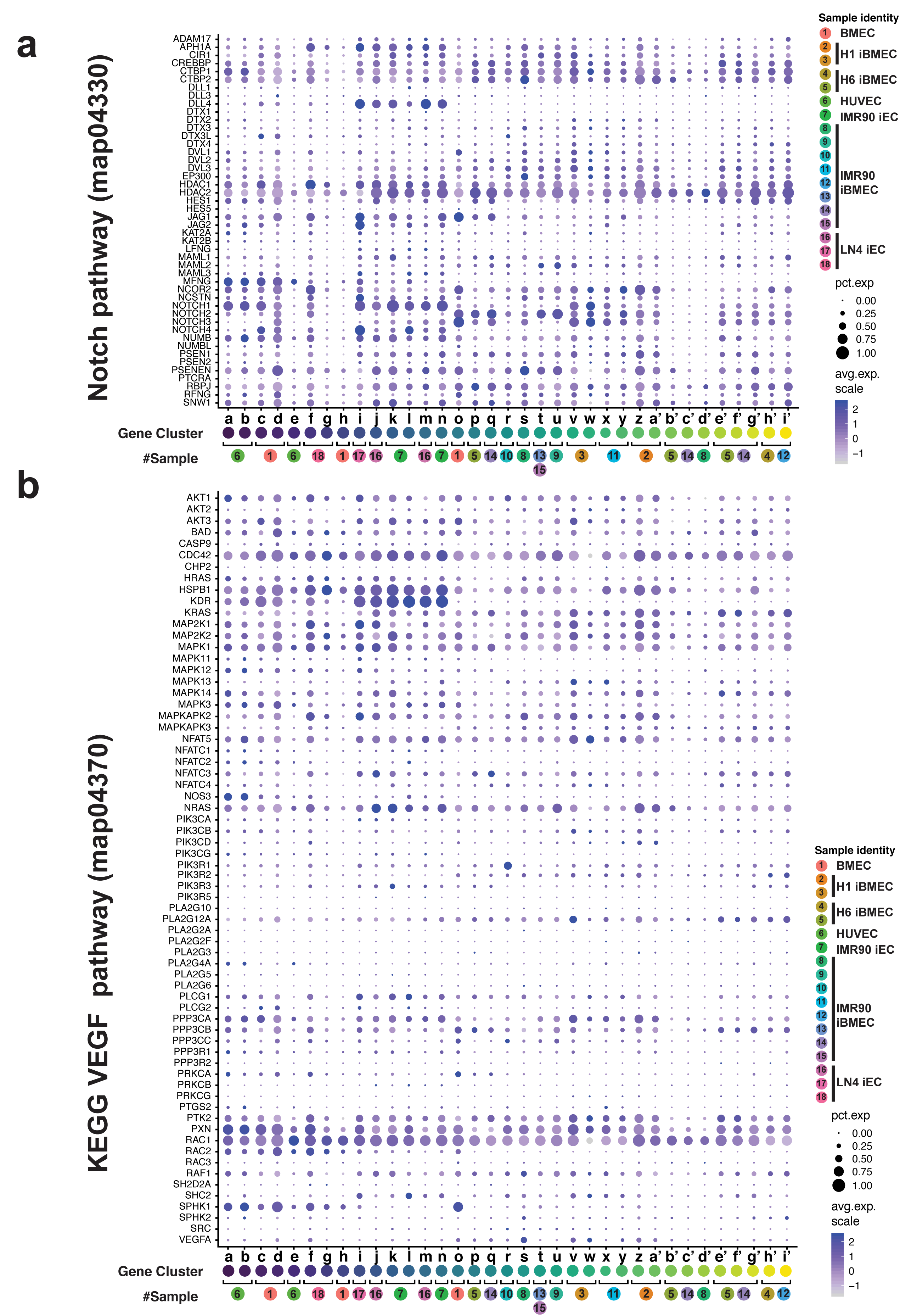
Expression of VEGF and NOTCH pathway genes across all samples. **a.** Dot plot comparing expression of Notch pathway genes (KEGG, map04330) across all single cell RNA sequencing derived cell clusters. **b.** Dot plot comparing expression of VEGF pathway genes (KEGG, map04370) across all single cell RNA sequencing derived cell clusters.

**Extended Data Figure 5.**
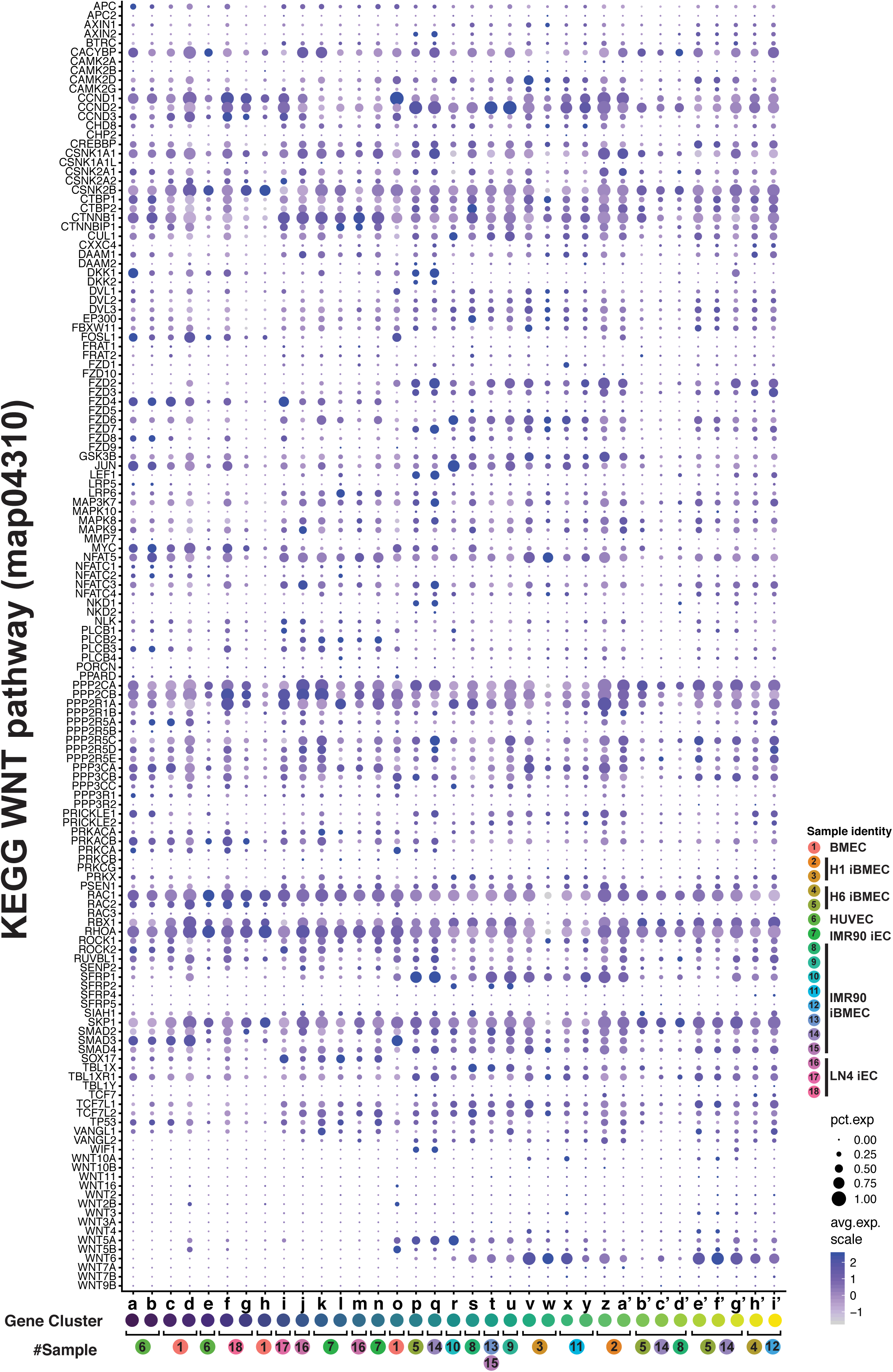
Expression of WNT pathway genes across all samples. Dot plot comparing expression of WNT pathway genes (KEGG, map04310) across all single cell RNA sequencing derived cell clusters.

**Extended Data Figure 6.**
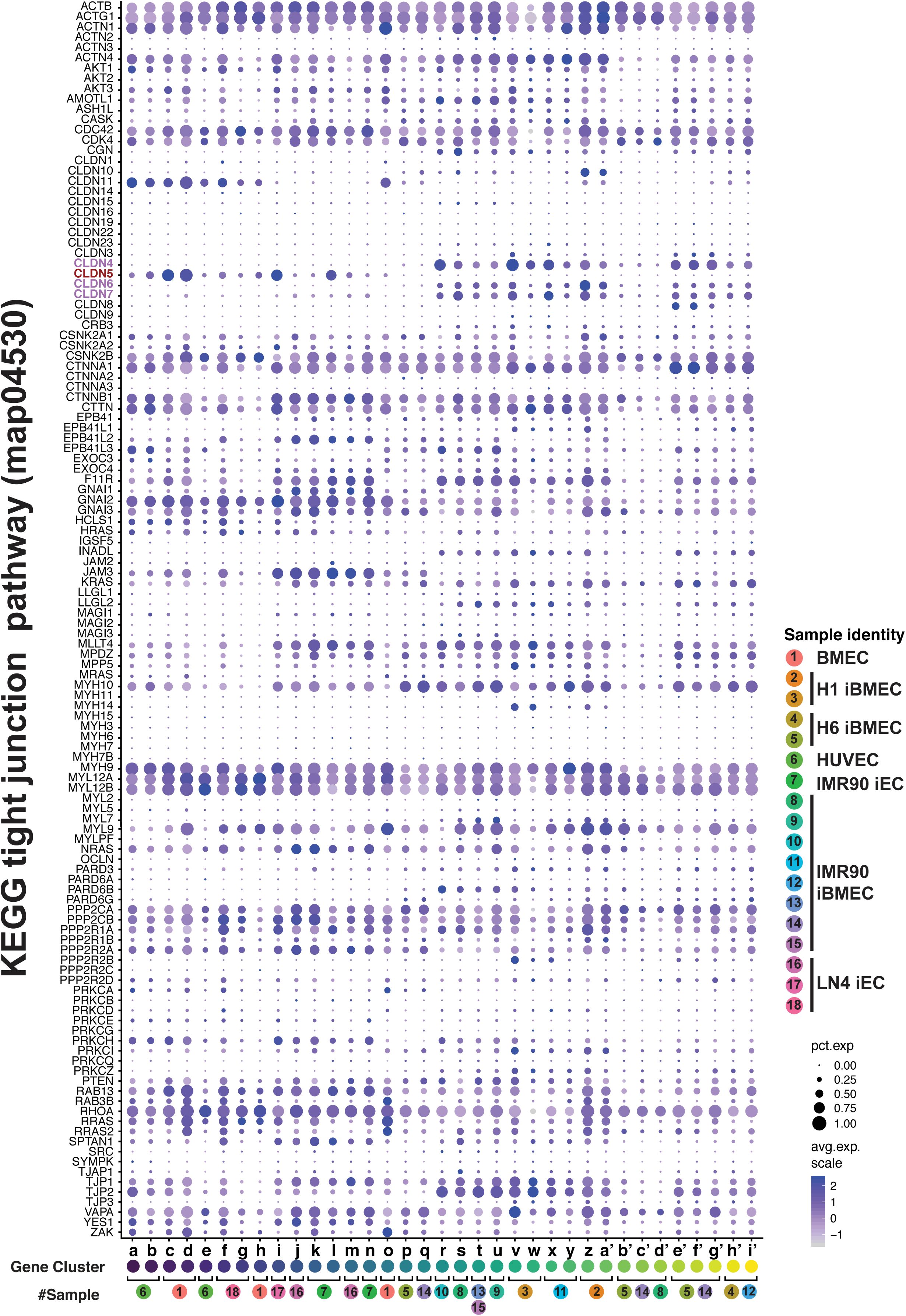
Expression of tight junction pathway genes across all samples. Dot plot comparing expression of tight junction pathway genes (KEGG, map04530) across all single cell RNA sequencing derived cell clusters. Genes highlighted show difference in Claudin expression profiles of iBMECs compared to iECs and primary ECs.

**Extended Data Figure 7.**
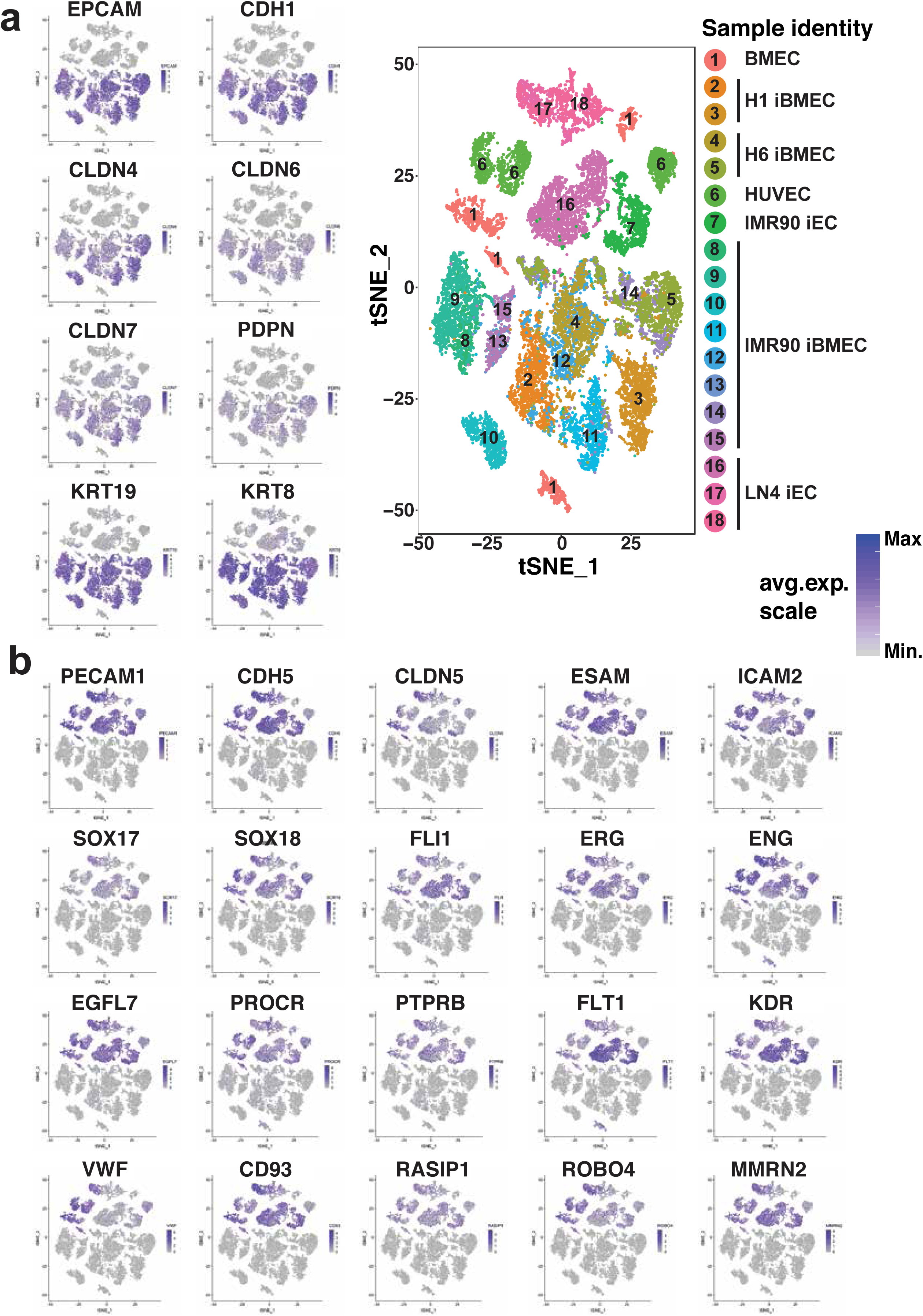
Distribution of endothelial and epithelial gene expression. **a.** scRNA expression profiles displayed on a t-distributed stochastic neighbor embedding (t-SNE) plots showing distribution of cells expression epithelial genes. **b.** scRNA expression profiles displayed on a t-distributed stochastic neighbor embedding (t-SNE) plots showing distribution of cells expression endothelial genes.

**Extended Data Figure 8.**
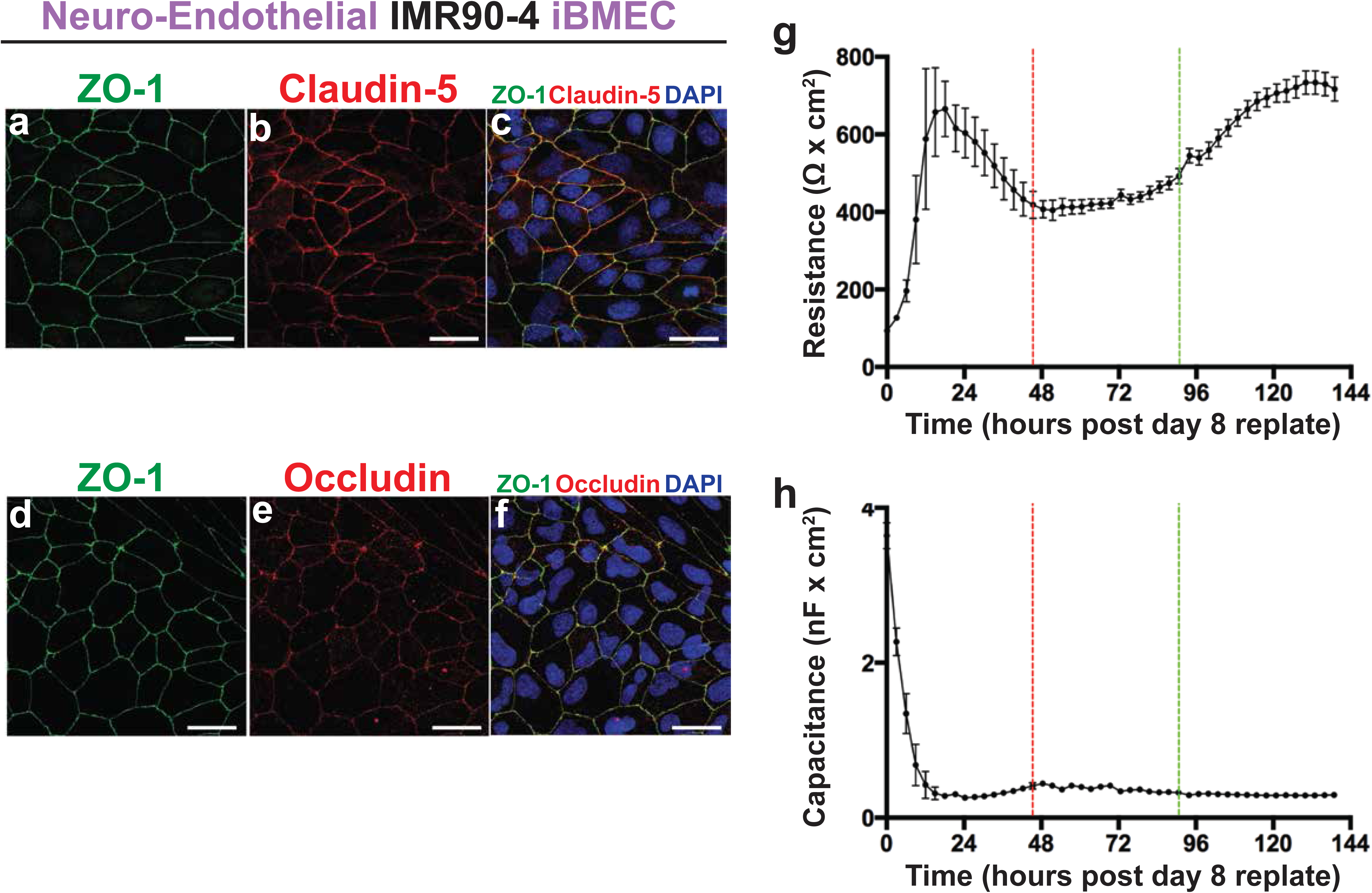
Immunofluorescence staining for tight junction proteins and TEER analysis of iBMECs generated with the neuroendothelial protocol. **a-f.** Immunofluorescence of CLAUDIN-5 (Red), OCCLUDIN (Red) and ZO-1 (Green) in IMR90-4 derived iBMECs using the published antibodies and DAPI (Blue). These antibodies recognize proteins at cell-cell junctions. Scale bars, 40µm. **g, h.** Measurements of Trans Endothelial Electrical Resistance and Capacitance of iBMECs over time. The cells were plated in ECIS plates after 8 days of neuroendothelial differentiation in a Collagen IV and Fibronectin substrate.

